# The Ciliopathy Gene *Ftm/Rpgrip1l* Controls Mouse Forebrain Patterning via Region-Specific Modulation of Hedgehog/Gli Signaling

**DOI:** 10.1101/408120

**Authors:** Abraham Andreu-Cervera, Isabelle Anselme, Alice Karam, Martin Catala, Sylvie Schneider-Maunoury

## Abstract

Primary cilia are essential for central nervous system development. In the mouse, they play a critical role in patterning the spinal cord and telencephalon via the regulation of Hedgehog/Gli signaling. However, despite the frequent disruption of this signaling pathway in human forebrain malformations, the role of primary cilia in forebrain morphogenesis has been little investigated outside the telencephalon. Here we studied development of the diencephalon, hypothalamus and eyes in mutant mice in which the *Ftm/Rgprip1l* ciliopathy gene is disrupted. At the end of gestation, *Ftm*^*-/-*^ fetuses displayed anophthalmia, a reduction of the ventral hypothalamus and a disorganization of diencephalic nuclei and axonal tracts. In *Ftm*^*-/-*^ embryos, we found that the ventral forebrain structures and the rostral thalamus were missing. Optic vesicles formed but lacked the optic cups. We analyzed the molecular causes of these defects. In *Ftm*^*-/-*^ embryos, *Sonic hedgehog* (*Shh*) expression was lost in the ventral forebrain but maintained in the zona limitans intrathalamica (ZLI), the mid-diencephalic organizer. In the diencephalon, Gli activity was dampened in regions adjacent to the *Shh*-expressing ZLI but displayed a higher Hh-independent ground level in the other regions. Our data uncover a complex role of cilia in development of the diencephalon, hypothalamus and eyes via the region-specific control of the ratio of activator and repressor forms of the Gli transcription factors. They call for a closer examination of forebrain defects in severe ciliopathies and for a search for ciliopathy genes as modifiers in other human conditions with forebrain defects.

## INTRODUCTION

The Hedgehog (Hh) pathway plays an essential role in forebrain patterning, as illustrated by its frequent perturbation in holoprosencephaly (HPE), a human condition defined as a defect in the formation of midline structures of the forebrain and face (Fernandes and Hébert, 2008; Muenke and Beachy, 2001). Null mouse mutants for *Shh* display a HPE phenotype (Chiang et al., 1996) and studies involving gene inactivation in mouse, lineage tracing, and loss- and gain-of-function approaches in chick identified multiple, successive functions of the Hh pathway in the diencephalon, hypothalamus and eyes (Furimsky and Wallace 2006; Vue et al., 2009; Jeong et al., 2011; Alvarez-Bolado et al., 2012; Haddad-Tovolli et al., 2012, 2015; Blaess et al. 2015; Zhang and Alvarez-Bolado 2016).

In vertebrates, transduction of Hh/Gli signaling depends on primary cilia, microtubular organelles with sensory functions. In the developing central nervous system, primary cilia are essential for proper dorso-ventral (DV) patterning of the spinal cord via modulating Hh signaling. Shh binds to its receptor Ptch1, which removes Ptch1 from the cilium and relieves the inhibition of the G-protein coupled receptor Smoothened (Smo) by Ptch1. Hh signaling at the cilium leads to the translocation of the Gli transcription factors into the nucleus and their activation into Gli activator form (GliA). In the absence of ligand, Gli2 and Gli3 are targeted to the proteasome in a cilium-dependent manner, giving rise to short forms with transcriptional repressor activity, among which Gli3R is a particularly strong repressor. Thus, the primary cilium is essential for the production of both GliR and GliA forms (Goetz and Anderson., 2010). In the forebrain, functional primary cilia are required for correct DV patterning of the telencephalon (Willaredt et al. 2008; Stottmann et al., 2009; Besse et al., 2011; Benadiba et al., 2012; Willaredt et al. 2013; Laclef et al., 2015) and for the proliferation of granule cell precursors in the dentate gyrus (Han et al., 2008). Surprisingly, despite the essential function of Hh signaling in the forebrain, the role of primary cilia outside the telencephalon has been little explored (Willaredt et al., 2013).

In this paper we study the function of the *Ftm/Rpgrip1l* gene in the forebrain. *RPGRIP1L* is a causal gene in severe human ciliopathies with brain abnormalities, Meckel-Gruber syndrome (MKS5 OMIM # 611561) and Joubert syndrome type B (JBTS7 OMIM # 611560) (Arts et al., 2007; Delous et al., 2007). The Rpgrip1l protein is enriched at the ciliary transition zone (TZ) and is essential for the TZ localization of many other ciliopathy proteins (Mahuzier et al., 2012; Reiter et al., 2012; Shi et al., 2017, Wiegering et al., 2018). Rpgrip1l is also required for proteasome activity at the cilium base and for autophagy (Gerhardt et al., 2015; Struchtrup et al., 2018).

*Ftm*^-/-^ mouse fetuses die at or shortly before birth with a ciliopathy phenotype (Delous et al., 2007; Vierkotten et al., 2007) and lack cilia in the developing telencephalon (Besse et al., 2011). Using this mutant, our lab has previously shown that primary cilia are required for telencephalic DV patterning. In *Ftm*^*-/-*^ embryos, the olfactory bulbs (OB) and corpus callosum (CC), two dorsal telencephalic structures, are missing, due to an expansion of the ventral telencephalon. The phenotype is rescued by introduction into the *Ftm* mutant of one allele of *Gli3*^*Δ699*^ (Besse et al., 2011; Laclef et al., 2015), which produces constitutively a short form of Gli3 with repressor activity (Hill et al., 2007). These studies demonstrate that the main role of cilia in telencephalic patterning is to permit Gli3R formation.

What is the role of primary cilia in other forebrain regions? Here we show that *Ftm*^*-/-*^fetuses display severely disorganized hypothalamus and diencephalon and lack eyes. Investigating the molecular causes of these defects, we find that *Shh* expression and Hh/Gli signaling are differentially affected in different forebrain regions. Our results uncover essential and diverse functions for Ftm/Rpgrip1l and cilia in Gli activity in patterning the forebrain and eyes.

## MATERIALS and METHODS

### Mice

All experimental procedures involving mice were made in agreement with the European Directive 2010/63/EU on the protection of animals used for scientific purposes, and the French application decree 2013-118. Mice were raised and maintained in the IBPS mouse facility, approved by the French Service for Animal Protection and Health, with the approval numbers C-75-05-24. The project itself has been approved by the local ethical committee “Comité d’éthique Charles Darwin”, under the authorization number 2015052909185846. *Gli3***^*Δ^699^*^** and *Ftm*-deficient mice were produced and genotyped as described previously (Böse et al., 2002; Besse et al., 2011). Mutant lines were maintained as heterozygous (*Ftm*^*+/****-***^ or *Gli3****^Δ699/+^***) and double heterozygous (*Ftm*^*+/****-***^; *Gli3****^Δ699/+^***) animals in the C57Bl6/J background. Note that the eye phenotype of the *Ftm*^*-/-*^ animals was totally penetrant in the C57Bl6/J background used here, unlike in C3H or mixed backgrounds (Delous et al., 2007 and C. Gerhardt, personal communication). The transgenic line Tg[GBS::GFP] was maintained in the C57Bl6/J background and genotyped as described (Balaskas et al., 2012). In analyses of *Ftm* mutant phenotypes, heterozygous and wild-type (wt) embryos did not show qualitative differences, and both were used as ‘control’ embryos. The sex of the embryos and fetuses was not analyzed. Embryonic day (E) 0.5 was defined as noon on the day of vaginal plug detection.

### Histology, In situ hybridization (ISH) and immunofluorescence (IF)

For whole-mount ISH, embryos were dissected in cold phosphate buffered saline (PBS) and fixed in 4% paraformaldehyde (PFA) in PBS for a time depending on the embryonic age and then processed as described in (Anselme et al., 2007). For histology and ISH on sections, embryos were dissected in cold PBS and fixed overnight in 60% ethanol, 30% formaldehyde and 10% acetic acid. Embryos were embedded in paraffin and sectioned (7 μm). Cresyl thionin staining and ISH were performed on serial sections, as described previously (Anselme et al., 2007, Besse et al., 2011, Laclef et al., 2015). For fluorescence ISH (FISH), immunodetection of the probe was done overnight at 4°C with anti-digoxigenin peroxidase-conjugated antibody (Roche), diluted 1/50 in maleate buffer supplemented with 2% Boehringer Blocking Reagent (Roche). Peroxidase activity was detected with FITC-coupled tyramide (1/50).

For IF, embryos were fixed overnight in 4% paraformaldehyde (PFA). E18.5 fetuses were perfused with 4% PFA. Immunofluorescence staining was performed on 14 **μ**m serial cryostat sections, as described previously (Anselme et al., 2007; Laclef et al., 2015), with antibodies against Shh (Cell signaling 2207, 1:200 and R&D Systems AF445, 1:200), Arl13b (Neuromab 75-287; 1:1500), FoxA2 (Abcam ab23630; 1:200), GFP (Aves GFP-1020; 1:200), Rpgrip1l (Besse et al., 2011; 1:800), Mash1 (BD Pharmigen 556604; 1:200). Secondary antibodies were Alexa-Fluor conjugates from Molecular Probes (1:1000). Nuclei were stained with DAPI (1:500).

### DiI/DiA labelling

Brains of E18.5 fetuses were dissected in PBS 1x and fixed overnight in 4% PFA. After three washes in PBS, were labelled by 1,1**′**-dioctadecyl-3,3,3**′**,3**′**-tetramethylindocarbocyanine perchlorate (DiI; Invitrogen D383) or 4-Di-16 ASP (4-(4- (Dihexadecylaminostyril)-N-Methylpyridinium iodide (DiA, Invitrogen D3883) crystals, in the cortex or in the diencephalon of control and *Ftm***^*-/-*^** brains, as indicated in Figure 1. Samples were kept for at least two weeks in PFA 4% at 37°C for the lipophilic dye to diffuse along the fixed cell membranes. Then, the brains were embedded in 4% agarose in PBS, and thick coronal vibratome (LeicaVT1000S) sections were made.

**Figure 1.**
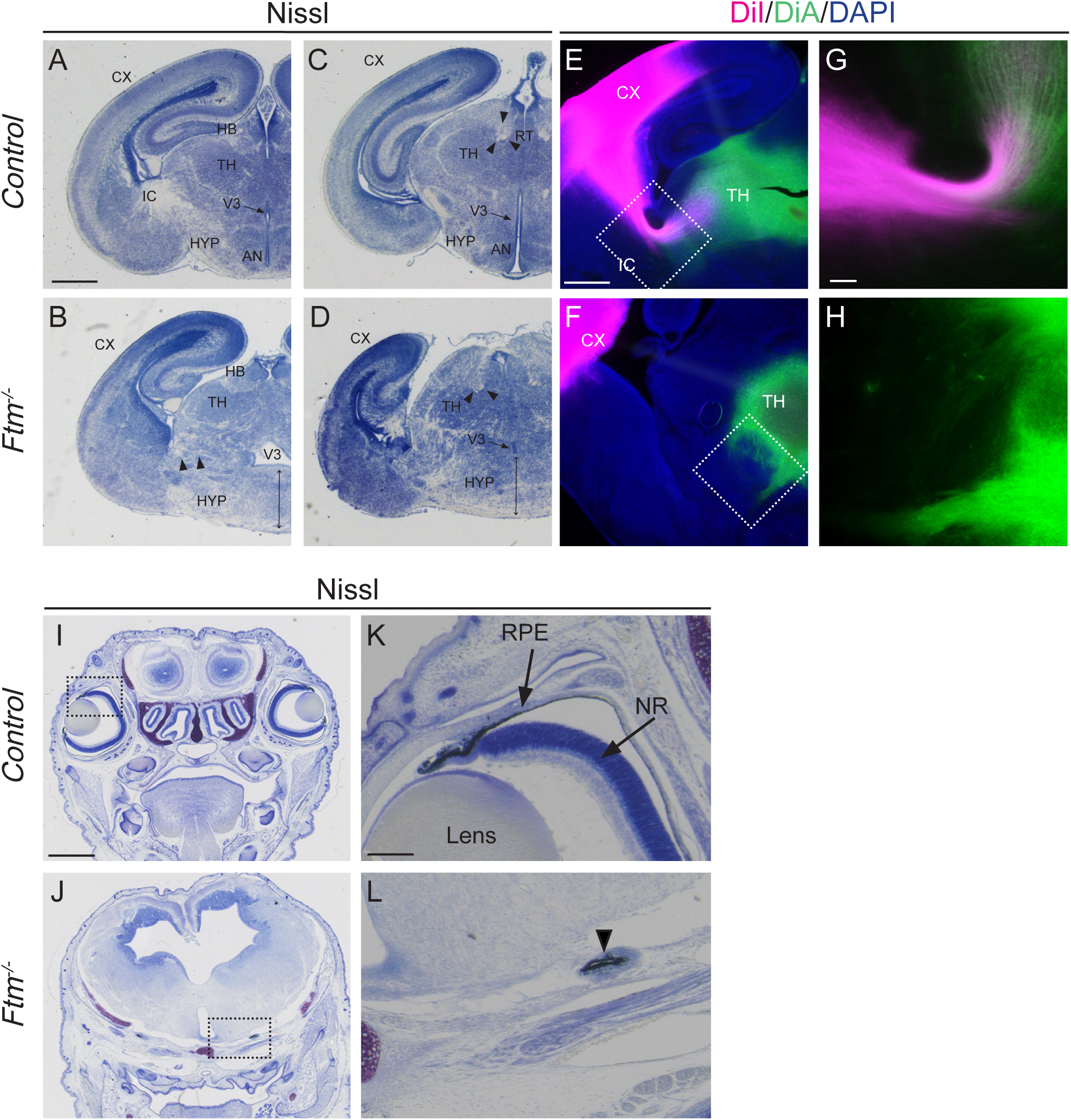
Histology and dye labelling of axon tracts in the brain of E18.5 fetuses. **A-D)** Nissl staining on coronal sections of the brain at two distinct antero-posterior levels of thalamic and hypothalamic regions in E18.5 wild type (A, C) and *Ftm*^*-/-*^ (B, D) fetuses. C, D are more posterior sections than A, B. Both levels of sections correspond to the ventral hypothalamus and the alar thalamus. Black arrowheads in B-D point to axon fascicles of the IC and Rt. Double black arrows in B, D point to the dysmorphic hypothalamus in *Ftm* mutants. **E-H)** Carbocyanine dye staining of corticothalamic (DiI, magenta) and thalamocortical (DiA, green) axons in E18.5 wild type (E, G) and *Ftm*^*-/-*^ (F, H) brains. G and H are higher magnification of the boxed regions in in E and F, respectively. **I-L)** Nissl staining on coronal sections at the level of the eyes of the head of E18.5 wild type (I, K) and *Ftm*^*-/-*^ (J, L) fetuses. K and L are higher magnification of the boxed regions in in I and J, respectively. The arrowhead in L points to remnants of the RPE. AN: anteroventral nucleus, C×: cortex; HB: Habenula; HYP: hypothalamus; IC: internal capsule; NR: neural retina; RPE: retinal pigmented epithelium; RT: retroflexus tract; TH: thalamus; V3 3^rd^ ventricle. Scale bars: 1 mm in A-D (shown in A) and in I and J (shown in I); 0.5 mm in E and F (shown in E); 0.1 mm in G and H (shown in G); 0.2 mm in K and L (shown in K).

### Image acquisition and quantification of fluorescence intensity

ISH images were acquired with a bright-field Leica MZ16 stereomicroscope. IF, FISH and axonal tract dye labelling images were observed with a fluorescent binocular (LeicaM165FC) and acquired with a confocal microscope (Leica TCS SP5 AOBS).

Fluorescence intensity was measured using the ImageJ software. For Shh-GFP immunofluorescence, adjacent squares of 50 μm side were drawn in the diencephalon, all along the ventricular surface from posterior to anterior. Total fluorescence intensity was measured in each square on three distinct optical sections. For each optical section, the background intensity was measured by taking three squares in the 3^rd^ ventricle, and the mean background intensity was subtracted from all the measurements of the same image. Images from three controls, three *Ftm*^*-/-*^ and two [*Ftm*^*-/-*^, *Gli3*^*Δ/+*^] embryos were used for quantification. For comparison, the measurements were aligned using as a reference the square corresponding to the AP level of the ZLI (point 12 of the abscissa on the diagrams in Figure 8K). The diagrams in Figure 8K indicate the mean intensity for each position of each genotype.

**Figure 8.**
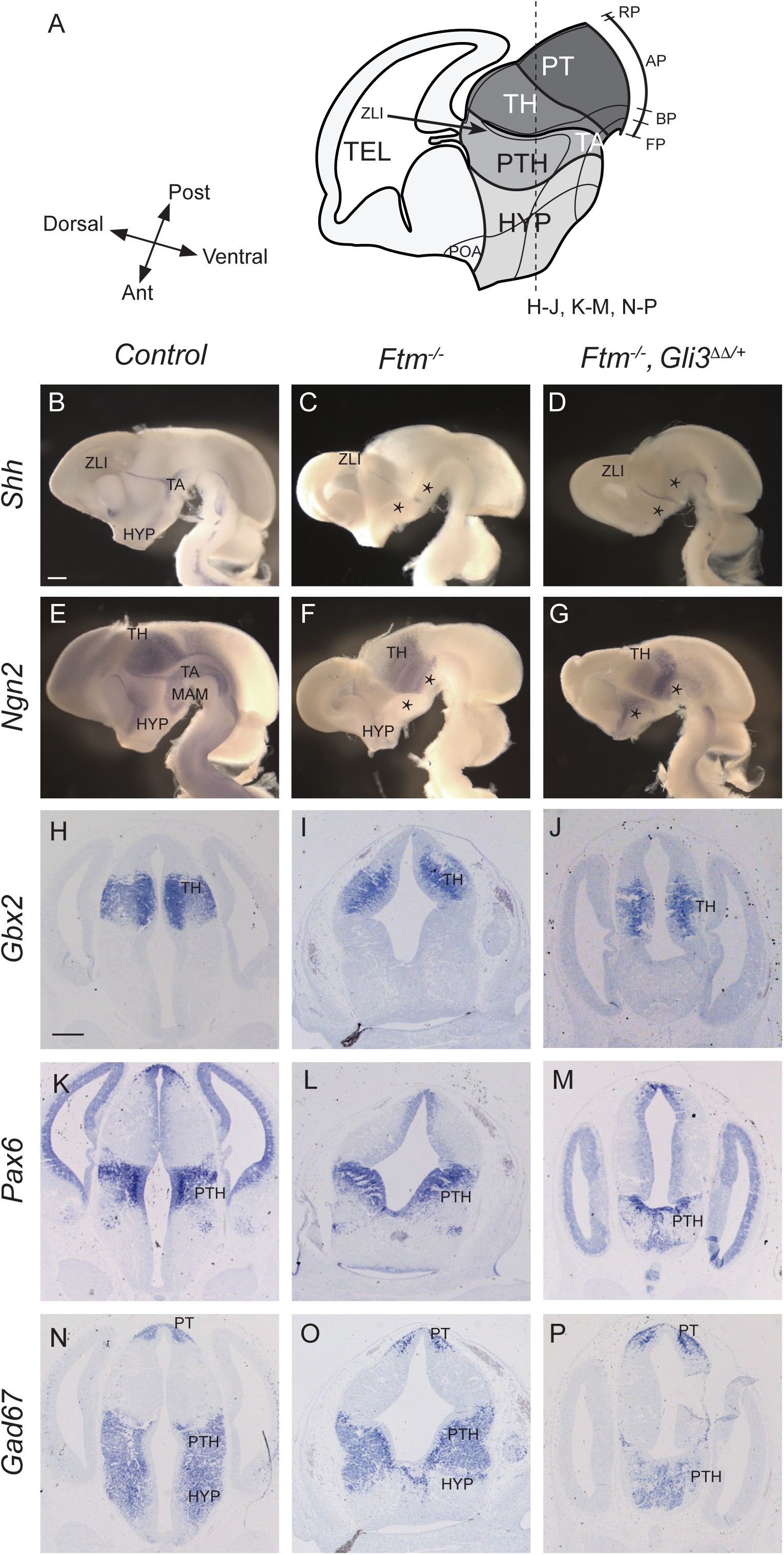
Diencephalon and hypothalamus patterning in compound [*Ftm, Gli3*^*Δ^699^*^] mutants. **A**) Schematic drawings of the E13.5 forebrain in sagittal view. The position of the coronal sections (H-P) shown below is indicated with dashed lines. **B-G**) Whole mount ISH with probes for *Shh* (B-D) or *Ngn2* (E-G) on sagittally-bisected brains viewed from the ventricular side. Black asterisks in C, D, F and G point to the absence of ventral forebrain. **H-P)** ISH on coronal sections with probe for *Gbx2* (H-J), *Pax6* (K-M) or *Gad67* (N-P). The genotype of the embryo is indicated on the top of the Figure. Ant: anterior; AP: alar plate; BP: basal plate; FP: floor plate; HYP: hypothalamus; RP, roof plate; Post: posterior; PT, pretectum; PTH, prethalamus; TA, tegmental areas; TEL: telencephalon; TH: thalamus; ZLI: zona limitans intrathalamica. Scale bars: 0.5 mm in all pictures (shown in B for coronal sections and in H for whole mount ISH).

For quantification of *Ptch1* FISH, adjacent squares of 20 μm side were drawn in the diencephalon, from posterior to anterior, at two apico-basal levels: along the ventricular surface and about 40 μm away from the ventricular surface. Total fluorescence intensity was measured in each square on three distinct optical sections. Images from 4 controls, 4 *Ftm*^*-/-*^ and 3 [*Ftm*^*-/-*^, *Gli3*^*Δ/+*^] embryos were used for quantification. For comparison, the measurements were aligned using as a reference the square corresponding to the AP level of the ZLI.

### Scanning electron microscopy

Embryos were dissected in 0.1 M sodium cacodylate (pH 7.4) and fixed overnight with 2% glutaraldehyde in 0.1 M sodium cacodylate (pH 7.4) at 4°C. Heads were then sectioned to separate the dorsal and ventral parts of the telencephalon, exposing their ventricular surfaces. Head samples were washed several times in 1.22 × PBS and post-fixed for 15 minutes in 1.22 × PBS containing 1% OsO4. Fixed samples were washed several times in ultrapure water, dehydrated with a graded series of ethanol and critical point dried (CPD 300, Leica) at 79?bar and 38?°C with liquid CO_2_ as the transition fluid and then depressurized slowly (0,025 bar/s). They were then mounted on aluminum mounts with conductive silver cement. Samples surfaces were coated with a 5 nm platinum layer using a sputtering device (ACE 600, Leica). Samples were observed under high vacuum conditions using a Field Emission Scanning Electron Microscope (Gemini 500, Zeiss) operating at 3 kV, with a 20 μm objective aperture diameter and a working distance around 3 mm. Secondary electrons were collected with an in-lens detector. Scan speed and line compensation integrations were adjusted during observation.

### Experimental design and statistical analysis

In all experiments, the number of embryos or fetuses analyzed was ≥ 3 for each genotype, unless otherwise stated. For the comparison of the number of cilia in the different diencephalic regions of control and *Ftm*^*-/-*^ embryos (Figure 9 P), quantification was made in 4 control and 4 mutant embryos. For statistical analysis, unpaired t-test was performed using the Prism software.

**Figure 9.**
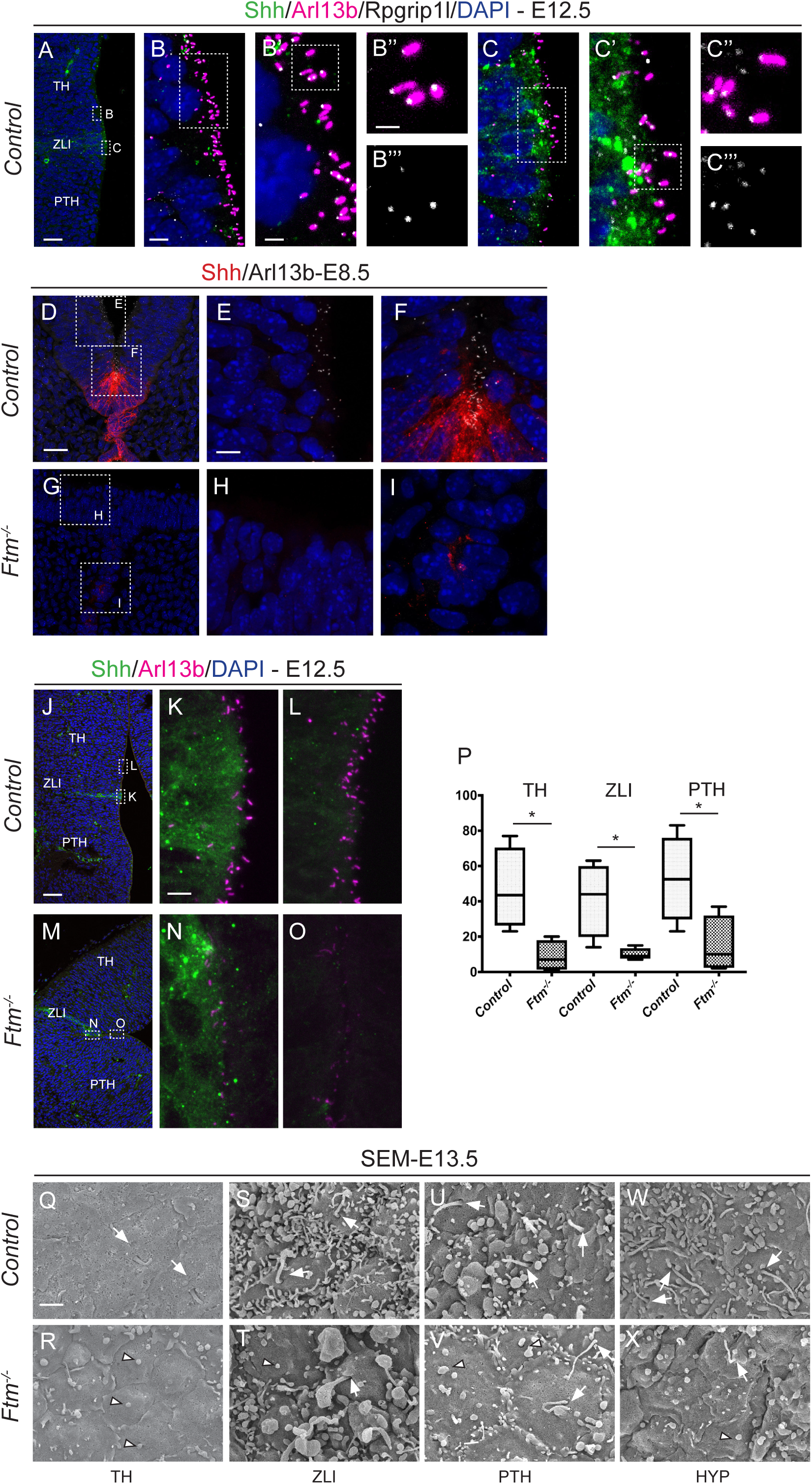
Cilia in the forebrain of *Ftm* mutants. **A-C’’’)** Immunofluorescence on coronal sections of E12.5 control embryos with antibodies for Shh (green), Arl13b (magenta) and Rpgrip1l (white). Nuclei are stained with DAPI. In B‴ and C‴, only Rpgrip1l is shown. **D-I)** Immunofluorescence on coronal sections of E8.5 control (D-F) and *Ftm*^*-/-*^ (G-I) embryos with antibodies for Shh (red) and Arl13b (white). Nuclei are stained with DAPI. White squares in D and G indicate the regions magnified in E, F and in H, I, respectively. **J-O)** Immunofluorescence on coronal sections of E12.5 control (J-L) and *Ftm*^*-/-*^ (M-O) embryos with antibodies for Shh (green) and Arl13b (magenta). In J and M, nuclei are stained with DAPI. White rectangles in J and M indicate the regions magnified in K, L and in N, O, respectively. **P)** Graph comparing the number of cilia in the diencephalon of control and *Ftm*^*-/-*^ embryos. * indicates that the statistical unpaired T-test is significant, with P value = 0.0223 for TH, 0.0263 for ZLI and 0.0421 for PTH. **Q-×)** SEM of the ventricular surface in different regions of control (Q, S, U, W) and *Ftm*^*-/-*^ (R, T, V, ×) hemisected brains. White arrows point to the base of cilia, white arrowheads point to button-like structures surrounded by a ciliary pocket, similar to those found in the cortex of *Ftm*^*-/-*^ embryos (Besse et al., 2011). Scale bars: 50 μm in A, J and M (shown in A and J); 20 μm in D, G (shown in D); 5 μm in B, C (shown in B) and in E, F, H and I (shown in E); 2 μm in B′ and C′ (shown in B′) and in K, L, N, O (shown in K); 1 μm in B″, B‴, C″, C‴ (shown in B″) and in Q-× (shown in Q).

## RESULTS

### *Ftm*^-/-^ fetuses at the end of gestation display microphthalmia and profound perturbations of the diencephalon and hypothalamus

Histological analysis combined with dye labelling of axonal tracts showed profound defects in the diencephalon and hypothalamus of *Ftm*^*-/-*^ fetuses at the end of gestation (E18.5; Figure 1). The ventral regions of the diencephalon and hypothalamus were particularly affected, with a highly dysmorphic ventral part and a perturbed position and shape of the 3^rd^ ventricle (Figure 1A-D). In wild type fetuses, habenular and thalamic nuclei were clearly visible in the dorsal region (Figure 1A, C). In *Ftm*^*-/-*^ fetuses, these nuclei were also present even if their organization was mildly perturbed (Figure 1B, D). In contrast, the ventral brain appeared highly disorganized in *Ftm*^*-/-*^ fetuses (Figure 1A-D). The ventral midline, normally thin in wild type, was enlarged in *Ftm*^*-/-*^, likely due to the absence of the most ventral region and secondary fusion of the lateral parts. The most medial hypothalamic nuclei (such as the anteroventral nuclei) were indistinguishable. The dorsal diencephalon and hypothalamus were present although malformed. In both regions, the axonal tracts (internal capsule (IC) and retroflexus tract (RT)) were defasciculated in *Ftm*^*-/-*^ brains (arrowheads in Figure 1B, D). Defects in cortico- thalamic (CTA) and thalamo-cortical (TCA) axonal tracts were confirmed with carbocyanine dye labelling (Figure 1E-H). In wild type brains, both CTA (magenta) and TCA (green) axons were visualized and colocalized in the IC (Figure 1E, G). In *Ftm*^*-/-*^ brains, neither CTA nor TCA grew sufficiently to reach the IC (Figure 1F, H). The eyes were absent in all *Ftm*^*-/-*^ fetuses (Figure 1I-L), only remnants of the retinal pigmented epithelium were observed under the brain (arrowhead in Figure 1L). We next focused on the developmental origin of these defects.

### Patterning of the diencephalon and hypothalamus is affected in *Ftm*^-/-^ embryos

The developing diencephalon is subdivided along the DV axis in roof, alar, basal and floor plates, and along the caudo-rostral axis in three regions or prosomeres, p1, p2 and p3. The alar plates of p1, p2 and p3 give rise to the pretectum (PT), thalamus (TH) and prethalamus (PTH), respectively (Figure 2A). The zona limitans intrathalamica (ZLI) is located at the junction between the TH and PTH. The ZLI acts as an organizer for the thalamus and prethalamus, regulating proliferation and cell fate in these two regions (Epstein, 2012; Hagemann and Scholpp, 2012; Zhang and Alvarez-Bolado, 2016).

**Figure 2.**
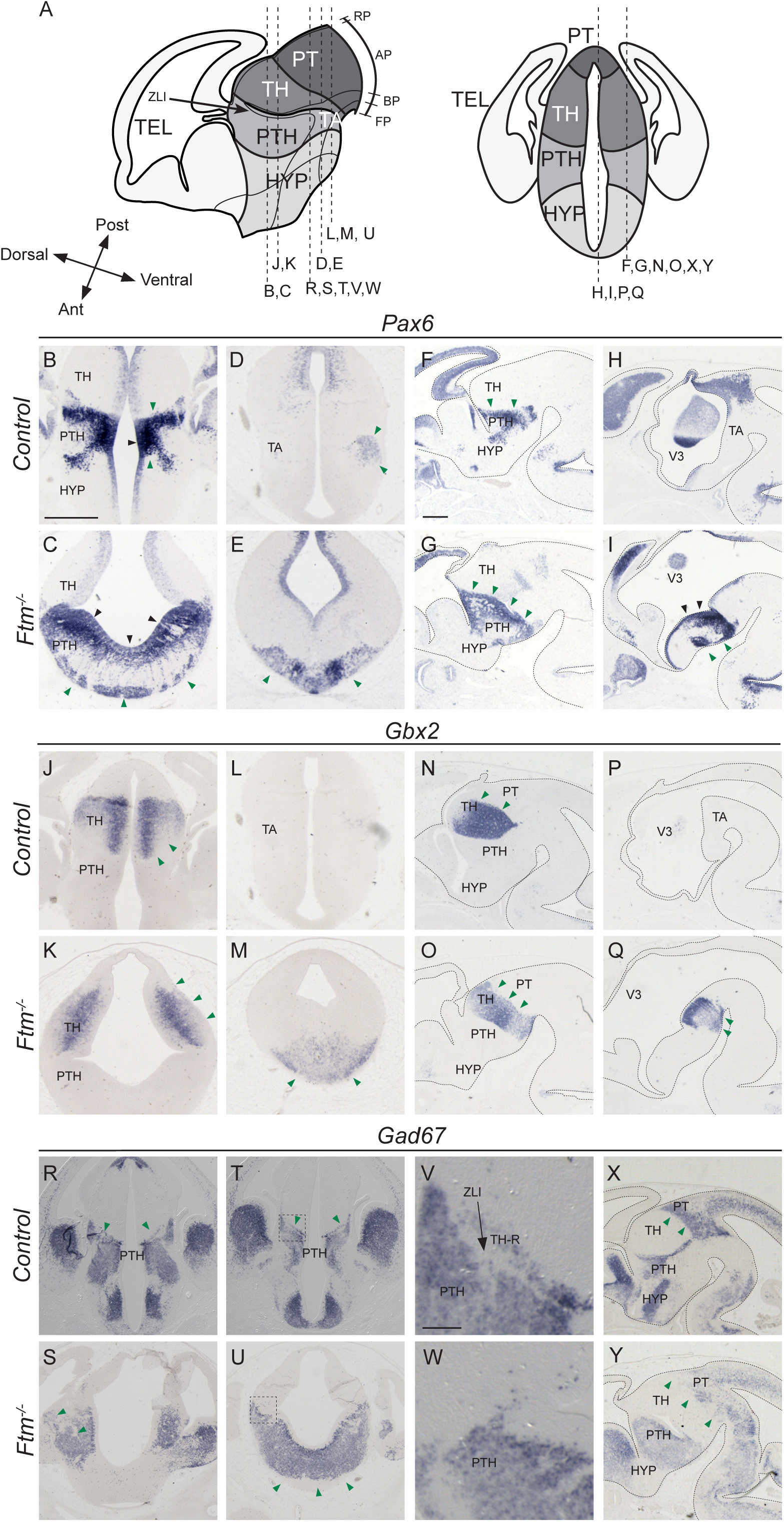
Diencephalon patterning at E13.5. **A)** Schematic drawings of the E13.5 forebrain in sagittal (left) and coronal (right) views. The position of the coronal (B-E, J-M, R-W) and sagittal (F-I, N-Q, ×, Y) sections shown below is indicated with dashed lines. Note that in the left diagram, antero-posterior and dorso-ventral axes are indicated at the level of the ZLI. **B-Y**) In situ hybridisation with probes for *Pax6* (B-I), *Gbx2* (J-Q) and *Gad67* (R-Y) in coronal sections at two distinct antero-posterior levels (B-E, J- M, R-W) and in sagittal sections at lateral (F, G, N, O, ×, Y) and medial (H, I, P, Q) levels. The genotype (*control* or *Ftm*^*-/-*^) is indicated on the left of the Figure, where control stands for *Ftm*^*+/+*^ or *Ftm*^*+/-*^. In sagittal sections, the brain is outlined with dotted lines. Black and green arrowheads point to neuronal progenitors and neurons, respectively. Ant: anterior; AP: alar plate; BP: basal plate; FP: floor plate; HYP: hypothalamus; RP: roof plate; Post: posterior; PT: pretectum; PTH: prethalamus; TA: tegmental areas; TEL: telencephalon; TH: thalamus; TH-R: rostral thalamus; ZLI: zona limitans intrathalamica; 3V: third ventricle. Scale bars: 0.5 mm in all pictures (shown in B for coronal sections and in F for sagittal sections) except in V and W where scale bar is 100 μm (shown in V).

To investigate diencephalon patterning in *Ftm* mutants, we performed *in situ* hybridization (ISH) for genes expressed in these different regions, on coronal and sagittal sections of E13.5 embryos. We first used the alar plate-expressed genes *Pax6* (PTH), *Gbx2* (TH) and *Gad67* (PTH and PT; Figure 2) encoding, respectively, two transcription factors involved at multiple steps of brain patterning and neurogenesis and a subunit of the glutamate decarboxylase involved in the synthesis of GABA (Stoykova and Gruss, 1994; Stoykova et al., 1996, Miyashita-Lin et al, 1999; Katarova et al., 2000; Hevner et al., 2002). We found that the expression domains of these genes were expanded along the DV axis in *Ftm*^*-/-*^ embryos (Figure 2). In control embryos, robust *Pax6* expression was detected in both the ventricular and subventricular zones (VZ and SVZ) of the PTH as well as in differentiating neuronal populations (Figure 2B, D, F, H). *Pax6* was also more faintly expressed in the VZ of the adjacent regions. In *Ftm*^*-/-*^ embryos, we observed a ventral expansion of the *Pax6* expression domain, which now reached the ventral midline (green arrowheads in Figure 2C, E, G, I). In addition, in anterior coronal sections, the hypothalamic, *Pax6*-negative region was absent from the sections shown (Figure 2E). *Gbx2* expression in control embryos was observed in differentiating neurons of the TH but not in the tegmental areas (TA) of the diencephalon (Figure 2J, L, N, P). In *Ftm*^*-/-*^ embryos, *Gbx2* expression expanded ventrally (Figure 2K, M, O, Q). *Gad67* expression in the control diencephalon was widespread in neurons of the PT and PTH and absent from diencephalic TA (Figure 2R, T, ×). In *Ftm*^*-/-*^ embryos, the PT and PTH expression domains expanded ventrally (Figure 2S, U, Y). The ventral expansion of the diencephalic alar plate and the reduction of the basal plate in *Ftm*^*-/-*^ embryos were confirmed using additional marker genes, *Ebf1* for PT, *Lhx2* for TH and *Six3* for PTH (Garel et al., 1997; Nakagawa and O’Leary, 2001; Puelles et al., 2006; Figure 3 B-E and data not shown).

**Figure 3.**
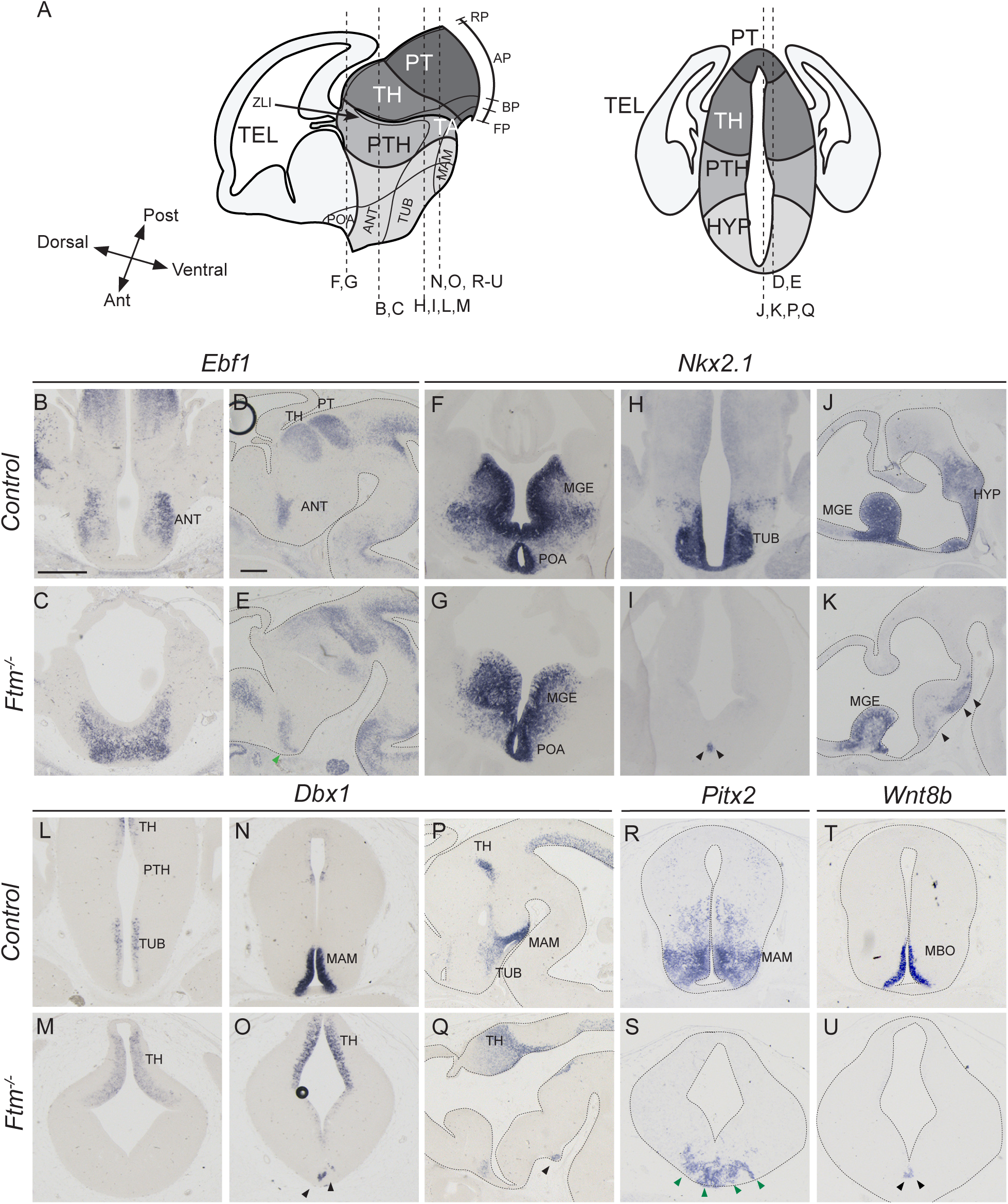
Hypothalamus patterning at E13.5. **A**) Schematic drawings of the E13.5 forebrain in sagittal (left) and coronal (right) views. The position of the coronal (B,C, F-I, L-O, R-U) and sagittal (D, E, J, K, P, Q) sections shown below is indicated with dashed lines. Note that in the left diagram, antero-posterior and dorso-ventral axes are indicated at the level of the hypothalamus. **B-U**) In situ hybridisation with probes for *Ebf1* (B-E), *Nkx2.1* (F-K), *Dbx1* (L-Q), *Pitx2* (R, S) and *Wnt8b* (T, U) in coronal sections at different antero-posterior levels and in sagittal sections. The genotype (*control* or *Ftm*^*-/-*^) is indicated on the left of the Figure. Black and green arrowheads point to neuronal progenitors and neurons, respectively. In sagittal sections and in coronal sections in R-U, the brain is outlined with dotted lines. Ant: anterior; ANT: anterior hypothalamus; AP: alar plate; BP: basal plate; FP: floor plate; HYP: hypothalamus; MAM: mammillary area; MBO: mammillary body; MGE: medial ganglionic eminence; POA: preoptic area; Post: posterior; RP: roof plate; PT: pretectum; PTH: prethalamus; TA: tegmental areas; TEL: telencephalon; TH: thalamus; TUB: tuberal hypothalamus; ZLI: zona limitans intrathalamica. Scale bars: 0.5 mm in all pictures (shown in B for coronal sections and in D for sagittal sections).

The hypothalamus can be subdivided into three main regions, the mammillary area (MAM), the tuberal hypothalamus (TUB) and the anterior hypothalamus (ANT). According to the revised prosomeric model (Puelles et al., 2012; Zhang and Alvarez-Bolado 2016), the MAM and TUB are in the basal plate of the hypothalamus while the ANT (also called alar hypothalamus) is in the alar plate (Figure 3A). The preoptic area (POA), formerly considered as a hypothalamic region, is actually part of the telencephalon. *Nkx2.1* is expressed in the hypothalamus in response to Hh signals from the underlying mesendoderm (Dale et al., 1997, Zhao et al., 2012, Blaess et al., 2015). *Nkx2.1* is also expressed in two telencephalic structures, the POA and medial ganglionic eminence (MGE; Figure 3F, H, J). In *Ftm*^*-/-*^ embryos, the *Nkx2.1* expression domain was preserved in the telencephalon (Figure 3F, G, J, K) but strongly reduced in the hypothalamus (Figure 3H-K). *Dbx1* expression in progenitors of the TUB (Figure 3L, M, P, Q) and MAM (Figure 3N-Q) regions was severely reduced as well, whereas it was maintained and even expanded in the thalamus (Figure 3L-Q). Analysis of *Pitx2* (Figure 3R, S) and *Wnt8b* (Figure 3T, U) expression confirmed the reduction in the surface of the MAM in *Ftm*^*-/-*^ embryos. *Ebf1* expression in the ANT was still present but fused at the midline (Figure 3B-E).

These data strongly suggest a severe reduction or loss of the basal plate and ventral midline of the forebrain in *Ftm*^*-/-*^ embryos. Conversely, the alar plate of the diencephalon appears expanded ventrally at all anteroposterior levels.

### The rostral thalamus is absent in *Ftm*^*-/-*^ embryos

We took advantage of the expression of two proneural genes, *Ngn2* and *Mash1/Ascl1*, expressed in distinct and complementary progenitor domains (Fode et al., 2000), to analyse diencephalic subdivisions with greater precision. *Ngn2* is expressed in progenitors of most of the TH, in the ZLI and in the tegmental areas (TA) of the diencephalon, in a domain in the POA and in the dorsal telencephalon (Fode et al., 2000; Vue et al., 2007; Figure 4B, D, F, H). *Mash1* is expressed in progenitors of the PTH, in the prospective rostral thalamus (TH-R, see below) and in different hypothalamic subdivisions (McNay et al. 2006; Vue et al., 2007; Kim et al., 2008; Figure 4J, L, N, P). *In Ftm*^*-/-*^ embryos, *Ngn1* expression was lost in the TA (empty arrowheads) and activated ectopically in a salt-and-pepper manner in regions adjacent to the telencephalon (black arrowheads), suggesting a perturbation of the telencephalic-diencephalic boundary (Figure 4C, E, G, I). *Mash1* was still expressed in the PTH and HYP (Figure 4K, M, O, Q), but very reduced caudally (in the MAM, black arrowheads in Q). The analysis of *Ngn2* and *Mash1* expression also revealed a thickening of the progenitor domains in the TH and PTH at E12.5-E13.5 (Figure 4B-Q), suggesting a delay in neurogenesis and/or an increased proliferation potential of forebrain progenitors in *Ftm*^*-/-*^ embryos.

**Figure 4.**
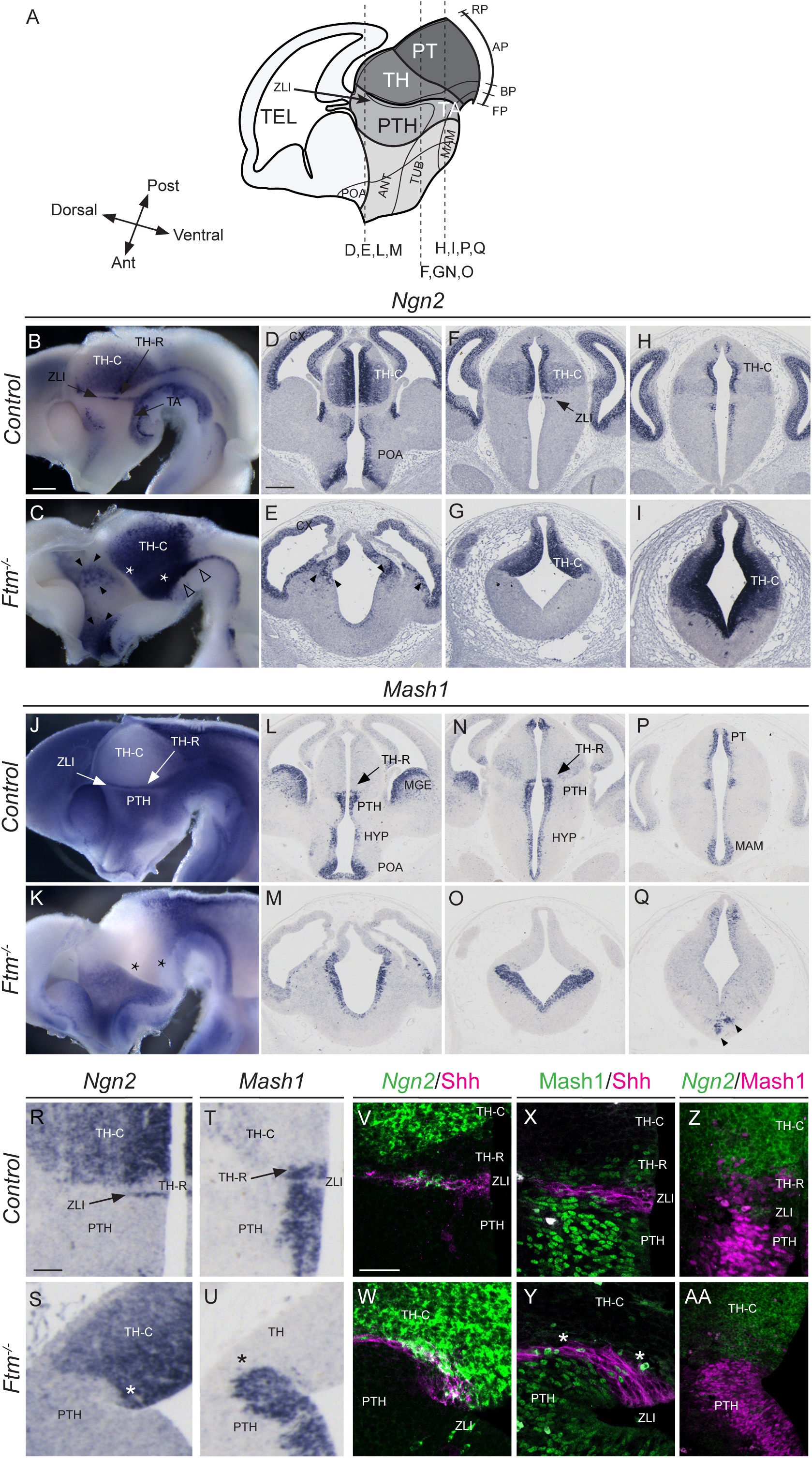
Progenitor domains at E12.5-E13.5. **A**) Schematic drawings of the E13.5 forebrain in sagittal view. The position of the coronal sections (D-I, L-Q) shown below is indicated with dashed lines. **B-U**) In situ hybridisation with probes for *Ngn2* (B-I, R, S) and Mash1 (J-Q, T, U) in whole mount hybridization on sagittally- bisected brains viewed from the ventricular side (B, C, J, K) or on coronal sections at different antero-posterior levels (D-I, L-Q, R-U). The genotype is indicated on the left of the Figure. **V, W)** Shh immunofluorescence (magenta) combined with *Ngn2* fluorescence in situ hybridization (green). **×, Y)** Double immunofluorescence for Shh (magenta) and Mash1 (green). **Z, AA)** Mash immunofluorescence (magenta) combined with Ngn2 fluorescence in situ hybridization (green). In C, black arrowheads point to patchy *Ngn2* expression in the prethalamus and white arrowheads point to missing *Ngn2* expression domain in the ventral forebrain. In P, black arrowheads point to remnants of the MAM. Asterisks in C, K, S and Y point to the absence of the *Ngn2*-negative, *Mash1*-positive TH-R in *Ftm*^*-/-*^ embryos. Ant: anterior; ANT: anterior hypothalamus; AP: alar plate; BP: basal plate; FP: floor plate; HYP: hypothalamus; MAM: mammillary area; POA: preoptic area; RP: roof plate; Post: posterior; PT: pretectum; PTH: prethalamus; TA: tegmental areas; TEL: telencephalon; TH: thalamus; TH-C: caudal thalamus; TH-R: rostral thalamus; TUB: tuberal hypothalamus; ZLI: zona limitans intrathalamica. Scale bars: 0.5 mm in B-Q (shown in B for whole mount ISH and in D for coronal sections); 100 μm in R-U (shown in R); 50 μm in V-Z and AA (shown in V).

The nested domains of *Ngn2* and *Mash1* expression in the diencephalon prefigure the intrinsic subdivision of the thalamus into anterior (TH-R) and posterior (TH-C) territories (Vue et al., 2007) (Figure 4B, J). *Ngn2* and *Mash1* domains in the TH and PTH, respectively, were continuous in *Ftm*^*-/-*^ embryos, suggesting a perturbation of thalamic subdivisions (Figure 4C, K, asterisks). This was confirmed by a closer examination of *Ngn2* and *Mash1* nested expression domains (Figure 4R-AA). We performed combined Shh/*Ngn2* (Figure 4V, W), Shh/Mash1 (Figure 4×, Y) and *Ngn2*/Mash1 (Figure 4Z, AA) fluorescence ISH and immunostaining to analyze the relationship of the different diencephalic domains with respect to the ZLI. In *Ftm*^*-/-*^ embryos, the domain of *Mash1* expression posterior to the Shh-positive ZLI was lost (white asterisks in Figure 4Y). The *Ngn2*-positive TH-C and Mash1-positive PTH domains abutted at the level of the ZLI (Figure 4W, Y, AA).

The TH-R contributes to GABAergic nuclei that participate in the subcortical visual shell, involved in the entrainment of the circadian rhythm (Delogu et al. 2012). Thus, neurons of the TH-R express *Gad67* like those of the PTH, while neurons of the TH-C do not (Figure 2V). In *Ftm*^*-/-*^ embryos, the stripe of *Gad67* expression in the thalamus was absent, confirming the loss of the TH-R (Figure 2V, W).

In conclusion, the TH-R is lost in *Ftm*^*-/-*^ embryos and the TH-C now abuts the ZLI. A diagram summarizing the *Ftm* mutant forebrain phenotype is provided in Figure 10A.

**Figure 10.**
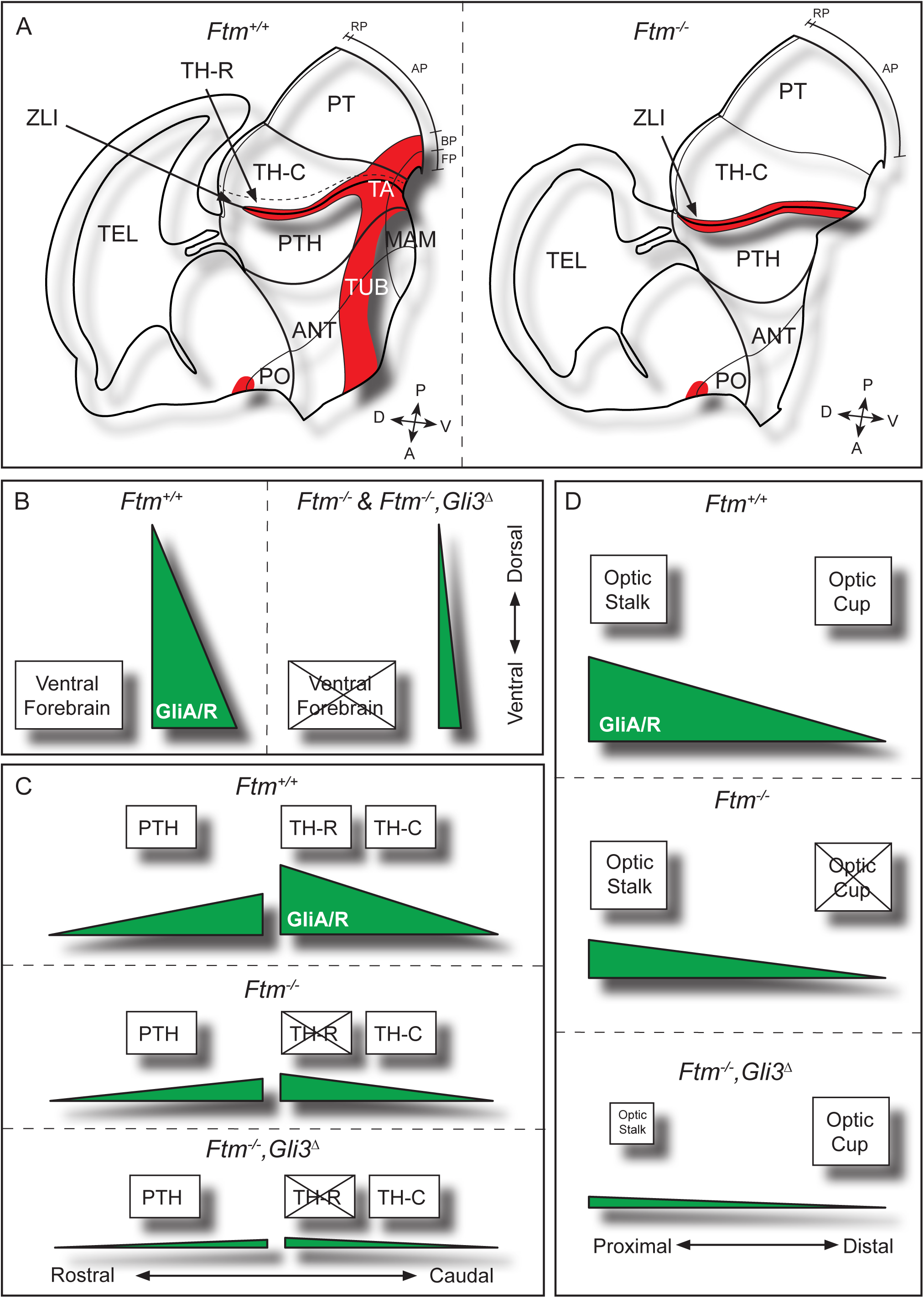
Schematics of forebrain patterning defects in *Ftm* embryos and their link to perturbations of Gli activity. **A)** Schematic drawings of the forebrain of E13.5 control (left) and *Ftm*^*-/-*^ (right) embryos. Shh expression domains are in red. **B-D)** Interpretive schematics of the GliA/GliR ratios (green) during ventral forebrain formation (B), alar diencephalon patterning (C) and optic vesicle patterning into optic stalk and optic cup (D) in control, *Ftm*^*-/-*^ and [*Ftm*^*-/-*^, *Gli3*^*Δ*^] embryos. **B)** A high GliA/GliR ratio is required for the formation of the ventral forebrain. In *Ftm*^*-/-*^ as well as in compound [*Ftm*^*-/-*^, *Gli3*^*Δ*^] embryos, the reduction this ratio causes a strong reduction of the ventral forebrain. **C)** In the alar diencephalon, a high GliA/GliR ratio is required for TH-R formation, while a lower ratio is sufficient for PTH and TH-C formation. In *Ftm*^*-/-*^ embryos the TH-R is lost but the ratio is sufficient for PTH and TH-C formation. **D)** Optic stalk formation requires a high GliA/GliR ratio, while the optic cup requires that only GliR is present. Low levels of GliA are sufficient for optic stalk formation in *Ftm*^*-/-*^ embryos. In contrast, the optic cup is not formed due to the reduction of GliR levels. In compound [*Ftm*^*-/-*^, *Gli3*^*Δ*^] embryos, the optic cup is rescued and the optic stalk is reduced (the eyes are closer to one another) due to the reintroduction of Gli3R. A : anterior; ANT : anterior hypothalamus; AP : alar plate; BP : basal plate; D : dorsal; FP : floor plate; MAM : mammillary hypothalamus; P : posterior; PT : pretectum; PTH : prethalamus; PO : preoptic area; RP : roof plate; TA : tegmental areas; TEL : telencephalon; TH-C : caudal thalamus; TH-R : rostral thalamus; TUB : tuberal hypothlamus; V : ventral; ZLI : zona limitans intrathalamica.

### Optic vesicles form in *Ftm*^*-/-*^ embryos and display patterning defects

Since eyes were absent in *Ftm*^*-/-*^ fetuses at the end of gestation (Figure 1I-L), we investigated eye formation and patterning at E11.5. Eye development begins with the formation of the eye field in the alar hypothalamus and its separation into two bilaterally symmetrical optic vesicles. The expanding optic vesicles induce the surface ectoderm to form the lens placodes. The optic vesicle separates into the optic stalk proximally and the optic cup distally. Then the optic cup invaginates with the lens placode, forming two layers, the outer layer differentiates into the retinal pigmented epithelium (RPE) and the inner layer into the neural retina (Furimsky and Wallace 2006).

We analyzed the expression patterns of the *Pax2, Vax2, Pax6* and *Chx10* transcription factor genes, which define distinct eye territories (Furimsky and Wallace 2006). At this stage, *Pax6* and *Pax2* are expressed in the optic cup and optic stalk, respectively (Figure 5F, K), where they repress each other. *Pax6* is required for optic cup formation, whereas *Pax2* null mice display increased optic cups at the expense of optic stalk (Schwarz et al., 2000). *Chx10* is also expressed in the optic cup (Figure 5P). *Vax2* is expressed in the ventral domain of the optic cup (Figure 5A), where it promotes ventral optic fates. In *Ftm*^*-/-*^ embryos, the neural retina was absent as assessed by the absence of *Chx10 and Vax2* expression (Figure 5B, Q). Only a tiny region of the RPE could be detected thanks to cell pigmentation (empty arrowheads in Figure 5B, G, L). *Pax2* was expressed, indicating the presence of the optic stalk (Figure 5G), which suggests correct eye field separation. Consistently, optic vesicles formed in E9 *Ftm*^*-/-*^ embryos as in controls (Figure 6S, U, V).

**Figure 5.**
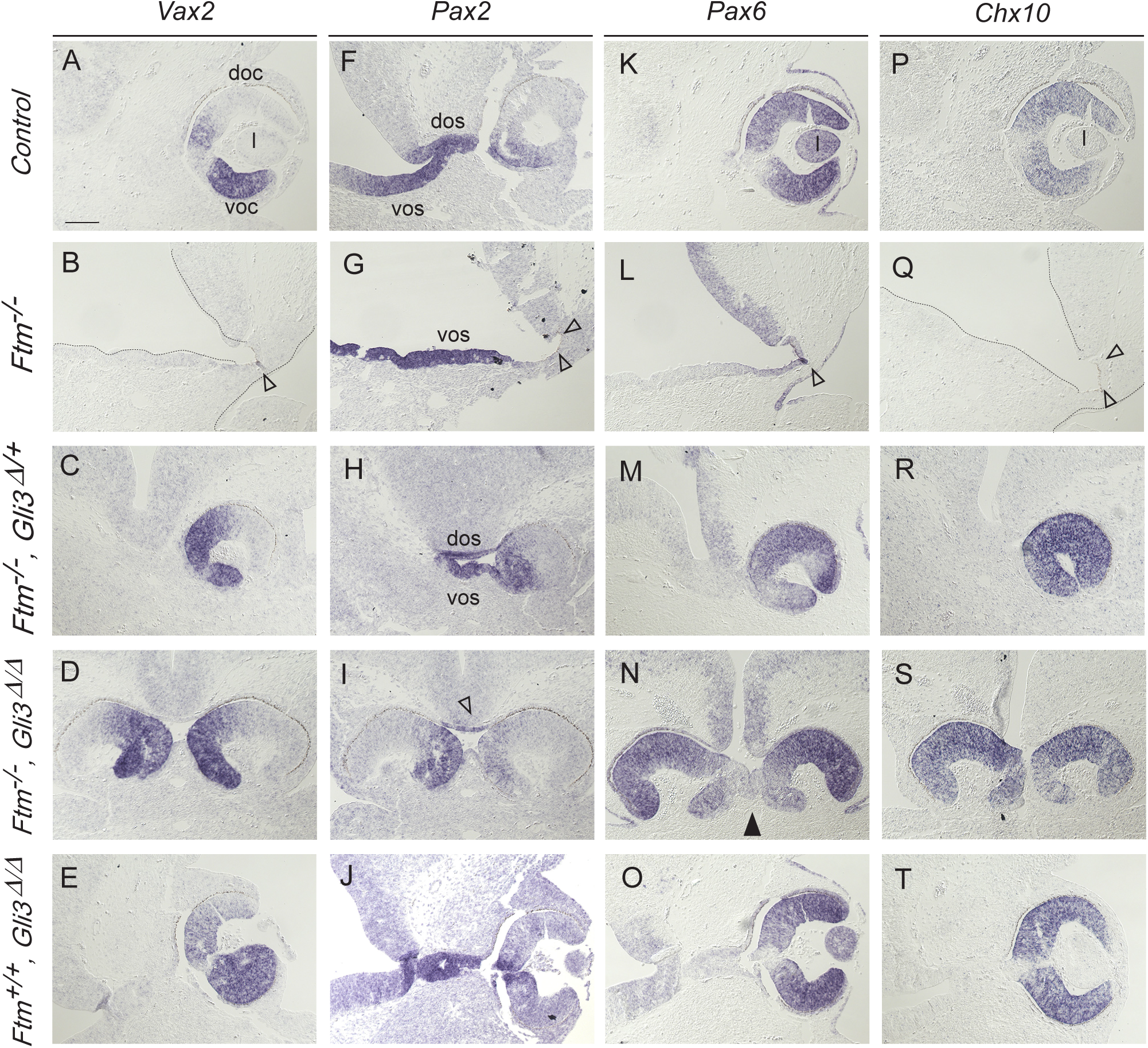
Eye morphogenesis in E11.5 embryos. **A-T)** ISH on coronal sections in the region of the eye of E11.5 *Ftm*^*+/+*^ (A, E, I, M), *Ftm*^*-/-*^ (B, F, J, N), *Ftm*^*-/-*^*, Gli3Δ/+* (C, G, K, O), *Ftm*^*-/-*^*, Gli3Δ/Δ* (D, H, L, P) and *Ftm*^*+/+*^*, Gli3Δ/Δ* (E, J, O, T) with probes for *Vax2* (A-E), *Pax2* (F-J), *Pax6* (K-O) and *Chx10* (P-T). Empty arrowheads in B, G, L and Q point to the missing optic cup; empty arrowhead in I points to the reduced optic stalk; black arrowhead in N points to partially fused optic cups. dos: dorsal optic stalk; l: lens; rpe: retinal pigmented epithelium; vos: ventral optic stalk. Scale bars: 100 μm (shown in A).

**Figure 6.**
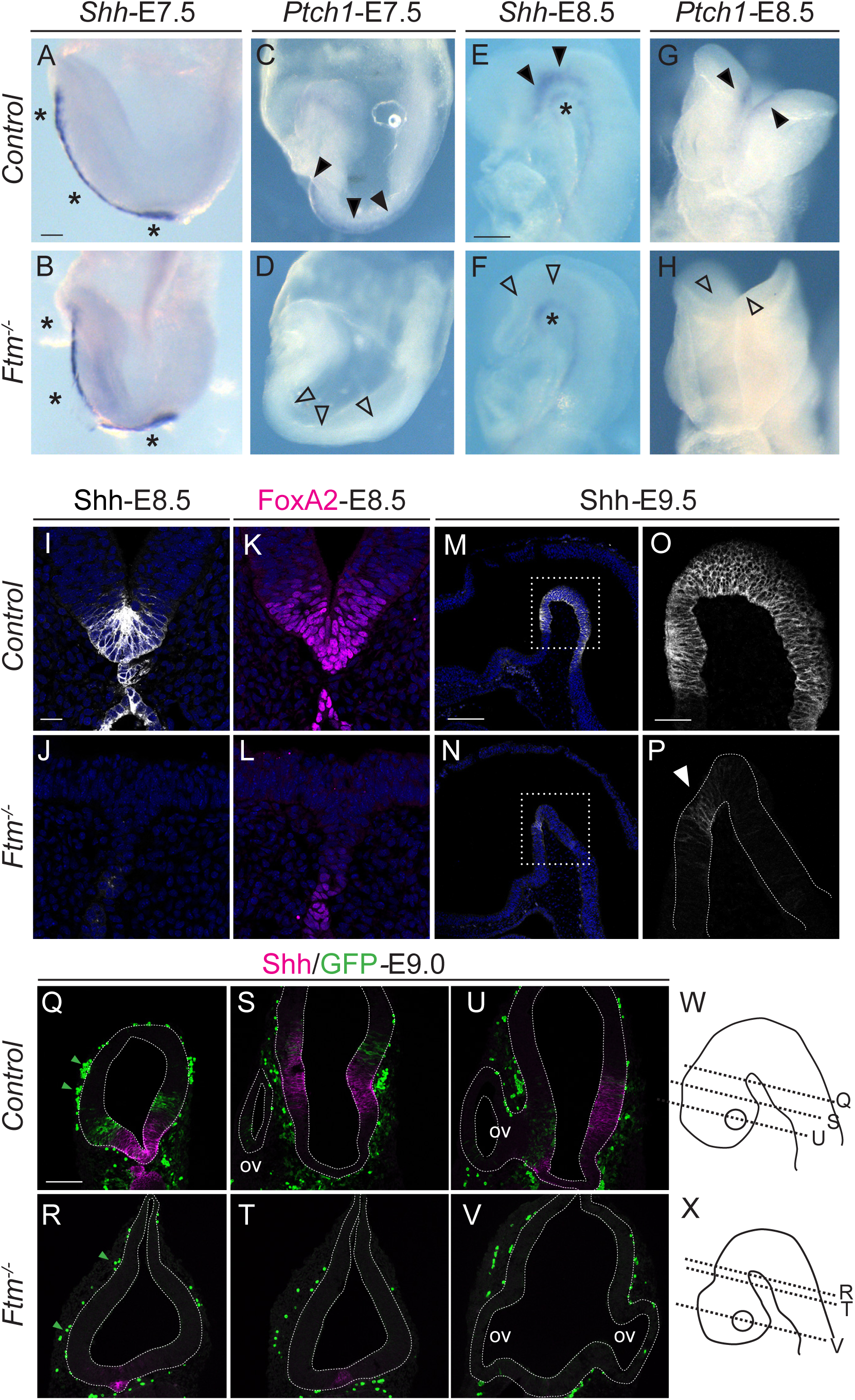
Hh expression and Hh/Gli signaling in the E7.5-E9.5 embryo forebrain. **A-H)** Whole-mount ISH on E7.5 (A-D) and E8.5 (E-H) embryos with probes for *Shh* (A, B, E, F) or *Ptch1* (C, D, G, H). Black arrowheads indicate *Shh* and *Ptch1* expression sites in the neural plate; black asterisks indicate *Shh* expression in the mesoderm underlying the neural plate. Empty arrowheads in D, F and H indicate absence of *Shh* and *Ptch1* expression in the neural plate of *Ftm*^*-/-*^ embryos. **I-P)** IF for Shh (I, J, M-P) and FoxA2 (K, L) on coronal sections of E8.5 (I-L) and sagittal sections of E9.5 (M-P) embryos. The white arrowhead in P points to the small dot of Shh expression in the basal plate of *Ftm*^*-/-*^ embryos. **Q-V)** Double IF for Shh and GFP in Tg[GBS::GFP] transgenic embryos. Green arrowheads point to GFP-positive blood cells. The genotypes are indicated on the left of the Figure. Control stands for *Ftm*^*+/+*^ or *Ftm*^*+/-*^. **W, ×)** Schematics indicating the approximate levels of sections in Q-V. Note that the sections are tilted, so they do not look bilaterally symmetric. Ov: optic vesicle. Scale bars: 50 μm in A- H (shown in A for A-D and in E for E-H); 20 μm in I-L (shown in I); 500 μm in M, N (shown in M); 100 μm in O, P (sown in O) and in Q-V (shown in Q).

In conclusion, in *Ftm*^*-/-*^ embryos, eye field separation occurs correctly but proximo-distal patterning of the optic vesicle is incorrect, leading to an absence of the optic cup and of lens induction.

### Hh expression and pathway activity are impaired in the forebrain of *Ftm*^*-/-*^ embryos

The reduction of the ventral forebrain in *Ftm* mutants suggests defects in the Hh pathway. To test this hypothesis, we analyzed Hh signaling activity in the forebrain by ISH and IF for *Shh* itself and for the Hh target genes *Ptc1, Gli1* and *FoxA2*. In addition, in order to obtain a context-independent assay of Hh transcriptional activity through Gli transcription factors binding to their DNA targets, we introduced into the *Ftm* mutant background the Tg[GBS::GFP] reporter transgenic line in which GFP expression is driven by a concatemer of Gli-binding sites (Balaskas et al., 2012).

In mouse embryos, *Shh* is initially expressed from E7.5 in axial tissues underlying the neural plate (notochord posteriorly and prechordal plate anteriorly), where it signals to the overlying neural plate to induce ventral structures. Hh signaling induces *Shh* expression in the ventral forebrain from E8.0 onwards (Dale et al., 1997). While *Shh* expression in the axial mesoderm was unperturbed in E7.5 *Ftm*^*-/-*^ embryos as compared to controls (asterisks in Figure 6A, B), *Ptch1* expression in the ventral neural plate (black arrowheads in Figure 6C) was lost in *Ftm*^*-/-*^ (empty arrowheads in Figure 6D). In E8.5 control embryos, *Shh* is still expressed in the mesendoderm underlying the brain (Figure 6E, I; asterisk in E). In addition, it is activated in the ventral neural tube, including the ventral forebrain (Dale et al., 1997; Alvarez-Bolado et al., 2012; Figure 6E, I, black arrowheads in E). In E8.5 *Ftm*^*-/-*^ embryos, *Shh* expression persisted in the notochord and prechordal plate (black asterisk in Figure 6F) but was not detected in the ventral neural tube and brain (Figure 6F, J, empty arrowheads in Figure 6F). *Ptch1* expression in two stripes surrounding the *Shh* expression domain in the ventral neural tube and brain of control embryos (black arrowheads in Figure 6G) was not found in *Ftm*^*-/-*^ (empty arrowheads in Figure 6H), consistent with the loss of *Shh* expression. *FoxA2*, a target of Hh signaling expressed in the ventral floor plate and in the ventral forebrain (Hallonet et al., 2002; Ribes et al., 2010; Figure 6K), was faintly expressed in the neural plate of *Ftm*^*-/-*^ embryos, confirming the very low Hh activity (Figure 6L).

At E9.0-E9.5, *Shh* expression is still present in the ventral diencephalon (Figure 6M, O, Q) but in the hypothalamus it is down-regulated in the most ventral region and activated in two lateral stripes in the basal plate (Figure 6S, U) (Szabo et al., 2009a; Alvarez-Bolado et al., 2012, Blaess et al., 2015). *Shh* expression was lost in the whole ventral forebrain of *Ftm*^*-/-*^ embryos, except in a tiny spot in the diencephalic ventral midline located roughly at the antero-posterior (AP) level of the future ZLI (Figure 6N, P, R, T, V). Gli activity assessed by Tg[GBS::GFP] was observed lateral to Shh expression domain in all forebrain regions of control embryos (Figure 6Q, S, U), whereas in *Ftm*^*-/-*^ embryos, Gli activity could not be detected (Figure 6R, T, V). Note that GFP is also expressed in blood cells and that this expression was still present in *Ftm*^*-/-*^ embryos (Balaskas et al., 2012; green arrowheads in Figure 6Q, R).

Overall our results show that Hh signaling activity is drastically reduced in the ventral forebrain of *Ftm*^*-/-*^ embryos as early as E8.5.

### *Shh* expression and Hh/Gli pathway activity show different perturbations in distinct domains of the E12.5 diencephalon

We next investigated Hh/Gli pathway activity at E12.5, when the ZLI is fully formed and secretes Shh to organize cell fate in the thalamus and prethalamus (Epstein, 2012; Zhang and Alvarez-Bolado, 2016). The ZLI was formed in both control and *Ftm*^*-/-*^ embryos, and its DV extent was increased in *Ftm*^*-/-*^ compared to control embryos (Figure 7A, B). In contrast, *Shh* expression was absent from the ventral forebrain of *Ftm*^*-/-*^ embryos (empty arrowheads in Figure 7B). In order to test whether Hh signaling was active at the ZLI, we analyzed *Ptch1* and *Gli1* expression as well as Gli transcriptional activity with Tg[GBS::GFP]. Surprisingly, *Gli1* and *Ptch1* were differently affected in *Ftm*^*-/-*^ embryos (Figure 7C-F). *Gli1* expression was dampened in the regions close to the *Shh* expression domains (Fig 7C, D). In contrast, *Ptch1* expression was totally downregulated in the thalamus and upregulated in the prethalamus and pretectum (Figure 7E, F). Using FISH on sections and signal quantification, we confirmed the differential expression of *Ptch1* on both sides of the ZLI in *Ftm*^*-/-*^ embryos and found that this upregulation was more striking at a distance from the ventricular surface (Figure 7M-N’, R, S). To test whether this reflected differential Gli activity on both sides of the ZLI, we observed GFP expression in Tg[GBS::GFP] embryos. At this stage, GFP-positive blood cells were present within the neural tube in all genotypes examined (green arrowheads in Figure 7G-I point to examples of these GFP-positive blood cells). We found that Gli activity was downregulated in the diencephalon and hypothalamus of *Ftm*^*-/-*^ embryos as compared to controls, in the ventral regions (Figure 7G, H) as well as on both sides of the ZLI (Figure 7J-K’’). However, Gli activity was not totally absent on both sides of the ZLI (Figure 7K-K’’). Moreover, in *Ftm*^*-/-*^ embryos, the Shh-positive ZLI appeared larger along the AP axis. Quantification of Shh and GFP immunofluorescence intensity confirmed the reduction of Gli activity and the increased width of the ZLI. In addition, it uncovered the presence in *Ftm* mutants of a ground level of Gli transcriptional activity along the whole AP extent of the TH and PTH, higher than in control embryos (Figure 7P, Q).

**Figure 7.**
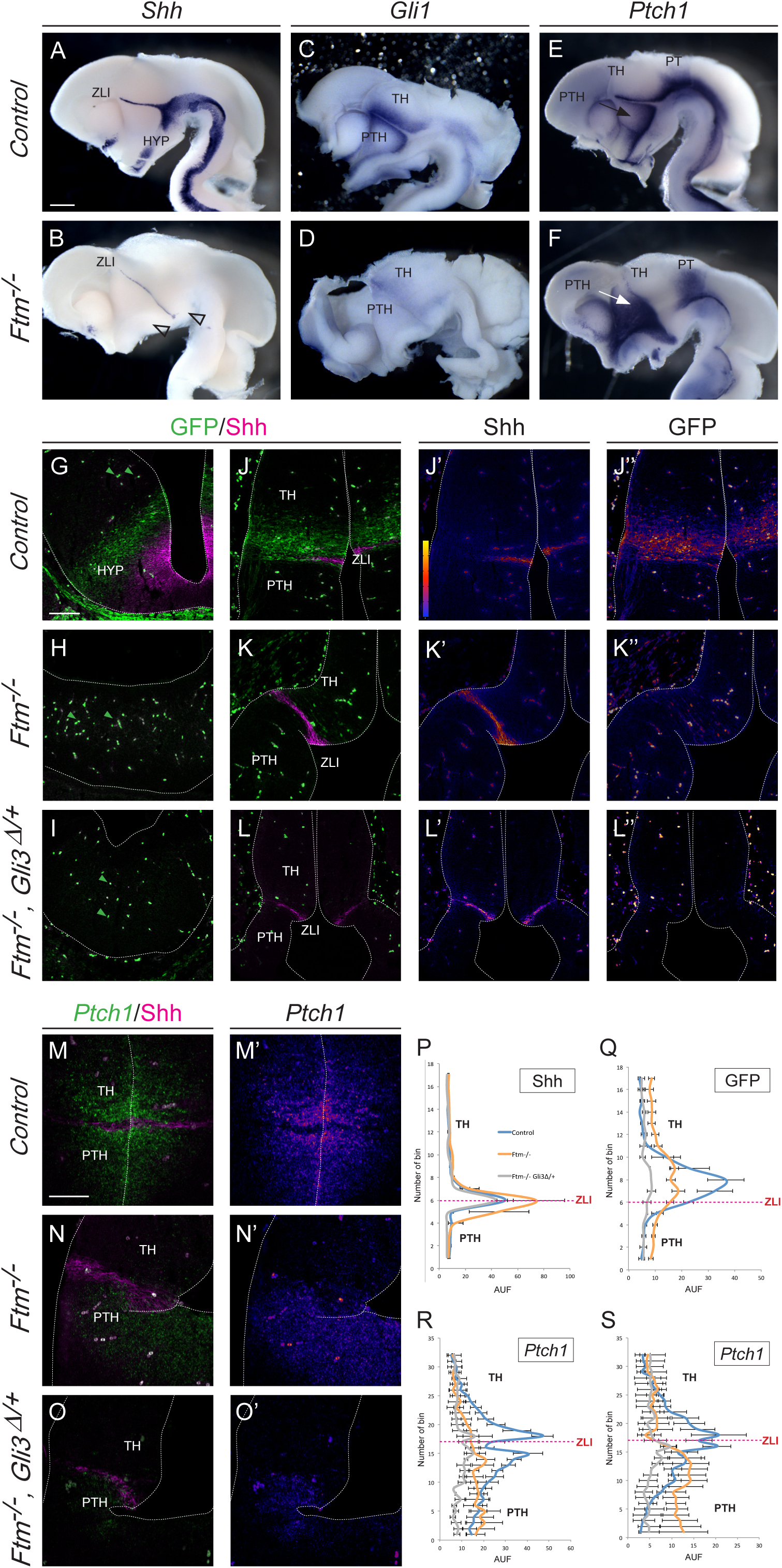
Hh expression and Hh/Gli signaling in the E12.5 embryo forebrain. **A-F)** Whole-mount ISH on E12.5 control (A, C, E) or *Ftm*^*-/-*^ (B, D, F) half-brains viewed from the ventricular surface, with probes for *Shh* (A, B), *Gli1* (C, D) or *Ptch1* (E, F). **G-L’’)** IF on coronal sections of control (G, J, J’, J’’), *Ftm*^*-/-*^ (H, K, K’, K’’) or [*Ftm, Gli3*^*Δ/+*^] (I, L, L’, L’’) Tg[GBS::GFP] embryos. IFs were performed with antibodies for Shh and GFP. In J’, J’’, K’, K’’, L’, L’’, fire versions of Shh and GFP are shown (fire scale in J’). Empty arrowheads in B point to the missing Shh expression domain in the ventral forebrain. **M-O’)** Combined fluorescence ISH *Ptch1* and IF for Shh on coronal sections for of the diencephalon of E12.5 control (M, M’), *Ftm*^*-/-*^ (N, N’) and compound [*Ftm*^*-/-*^*, Gli3*^*Δ/+*^] (O, O’) embryos. M’-O’ show fire versions of *Ptch1* FISH. Green arrowheads in. M, N and O point to GFP-positive blood cells. **P-S)** Diagrams showing the quantification of the intensity of Shh (P) or GFP (Q) IF and *Ptch1 FISH* (R, S) along the diencephalon. *Ptch1* FISH intensity was quantified next to the ventricular surface (R) or 40 μm away from the ventricular surface (S). Numbers on the abscissa relate to the position of the squares of quantification. Fluorescence intensity in ordinate is given in arbitrary units (AUF). HYP: hypothalamus; PT: pretectum; PTH: prethalamus; TH: thalamus; ZLI: zona limitans intrathalamica. Scale bars: 0.5 mm in A-F (shown in A); 100 μm in G-O’ (shown in G).

In conclusion, in the *Ftm*^*-/-*^ embryos, *Shh* expression is strongly reduced in the ventral forebrain but maintained and even expanded in the ZLI. Gli activity is dampened in regions adjacent to Shh-expressing domains and displays a ground level higher than in control embryos in other regions. The loss of Ftm also uncovered a differential prepattern of *Ptch1* and *Gli1* expression in different diencephalic prosomeres.

### Reintroduction of Gli3R into the *Ftm* background rescues aspects of the forebrain phenotype

The increased ground level of Gli activity in *Ftm*^*-/-*^ embryos is likely caused by the impaired production of Gli3R (Vierkotten et al., 2007, Besse et al., 2011). We thus tested how Gli activity in the diencephalon was modified in compound [*Ftm*^*-/-*^; *Gli3Δ/+*] embryos, by performing quantification of *GFP* and *Shh* expression in [*Ftm*^*-/-*^*; Gli3*^*Δ/+*^] mutant embryos harbouring Tg[GBS::GFP]. The *Gli3*^*Δ^699^*^ allele produces constitutively a short form of Gli3 with partial repressor activity (Hill et al., 2007; Cao et al., 2013). We found that both the ground level of Gli activity and the Shh-dependent Gli activity adjacent to the ZLI were reduced in these compound mutants (Figure 7I, L-L’’, P, Q). *Ptch1* expression in the prethalamus was also downregulated (Figure 7O, O’, R, S). Moreover, the increased width of *Shh* expression in the ZLI was rescued in double mutants (Figure 7J-O).

We then tested the consequences of Gli3R reintroduction on forebrain patterning and eye formation. ISH for *Shh, Ngn2, Gbx2, Pax6* and *Gad67* indicated that the reduction of the ventral forebrain was still observed and even worsened in [*Ftm*^*-/-;*^ *Gli3Δ/+*] (Figure 8B-P). As in *Ftm*^*-/-*^ embryos, the alar plate of the diencephalon was expanded ventrally (Figure 8E-P), and *Shh* expression was absent from the ventral forebrain but present in the ZLI (Figure 8C, D). In contrast, optic cup formation was totally restored in compound mutants, and the optic cup showed correct DV patterning (Figure 5C, D, H, I, M, N, R, S). However, the eyes were internalized and brought together in [*Ftm*^*-/-*^*; Gli3*^*Δ/+*^] embryos, and even more [*Ftm*^*-/-*^*; Gli3*^*Δ/Δ*^] embryos, and this was associated with a very reduced optic stalk (Figure 5C, D, H, I, M, N, R, S). We also analysed [*Ftm*^*+/+*^; *Gli3*^*Δ/Δ*^] embryos, which looked similar to controls (Figure 5E, J, O, T) as found in another study (Christoph Gerhardt, personal communication), indicating that only GliR is required for optic cup formation.

In conclusion, reintroducing Gli3R into the *Ftm* background rescues some of the defects of *Ftm*^*-/-*^ embryos, such as optic cup agenesis and ZLI enlargement, but not others such as the reduction of the forebrain basal plate and of the rostral thalamus. Moreover, it triggers optic stalk hypoplasia.

### Cilia of forebrain neural progenitors are severely reduced in number and malformed in *Ftm*^*-/-*^ embryos

In the telencephalon of *Ftm*^*-/-*^ embryos, neural progenitors are devoid of primary cilia (Besse et al., 2007). Since our data indicate that Hh activity is not totally lost in the diencephalon and hypothalamus of *Ftm*^*-/-*^ embryos, we tested the status of cilia in this region. We first analyzed Rpgrip1l expression in E12.5 controls and found that it was present at the ciliary transition zone in different diencephalic domains including the ZLI (Figure 9A-C’’’). We then compared cilia in the control and *Ftm* mutant brain at different stages by immunofluorescence for Arl13b. Cilia were present in the forebrain of E8.5 control embryos (Figure 9D-F) but were not detected in *Ftm*^*-/-*^ embryos (Figure 9G-I). In the E12.5 diencephalon, cilia were present in the TH, PTH and ZLI in control embryos (Figure 9J-L) and severely reduced in number in *Ftm*^*-/-*^ embryos (Figure 9M-O and P). Arl13b staining was less intense in the remaining cilia (Figure 9M-O). To analyze cilia shape in greater detail we performed scanning electron microscopy (SEM) of the ventricular surface of E13.5 control and *Ftm*^*-/-*^ brains, at different AP levels: in the TH, ZLI, PTH and HYP (Figure 9Q-×). In control embryos, cilia of about 1 μm in length were found in the TH, PTH and ZLI (arrows in Figure 9Q, S, U), while in the HYP cilia were frequently up to 3 μm long (arrows in Figure 9W). Cilia were more difficult to recognize in the ZLI since the ventricular surface of the cells was rich in protrusions and vesicles (Figure 9S). In the diencephalon and hypothalamus of the *Ftm*^*-/-*^ forebrain, cilia were in majority reduced to button-like structures (arrowheads in Figure 9R, T, V, ×), with a few very long cilia often abnormal in shape (arrows in Figure 9R, T, V, ×). These remaining cilia were present in all regions, but more frequently in the ZLI (Figure 9T).

In conclusion, cilia were absent from the forebrain of *Ftm*^*-/-*^ embryos as soon as E8.5. At E12.5 they were reduced in number in the diencephalon and hypothalamus, and the remaining cilia were longer than in controls and often presented an abnormal shape.

## DISCUSSION

The role of cilia in the forebrain has been little studied outside of the telencephalon. In this paper we have studied the role of the *Ftm/Rpgrip1l* ciliopathy gene in patterning of the diencephalon, hypothalamus and eyes. At the end of gestation, *Ftm*^*-/-*^ fetuses displayed anophthalmia, reduction of the ventral hypothalamus and disorganization of diencephalic nuclei and axonal tracts. We examined the developmental defects underlying this phenotype. *Ftm*^*-/-*^ embryos showed a severe reduction of ventral forebrain structures accompanied by a dorsoventral expansion of alar diencephalic domains and a loss of the rostral thalamus (Figure 10A). Optic vesicles formed but optic cup morphogenesis did not occur. Investigating the Hh pathway, we identified region-specific perturbations of Gli target gene expression and of Gli activity. Combined with our previous studies (Besse et al., 2011; Laclef et al., 2015), our data lead to a global understanding of the role of primary cilia in forebrain patterning and morphogenesis of their relationship with Hh signaling.

Do the forebrain defects of *Ftm* mutants correspond to a ciliary phenotype? Apart from the disorganization of the diencephalic-telencephalic boundary (Willaredt et al., 2008; 2013), the defects observed in this study have not been reported in other ciliary mutants. It was thus important to study the number and integrity of cilia in the diencephalon and hypothalamus of *Ftm*^*-/-*^ embryos at different stages. We found a near-total loss of cilia in the progenitors of the forebrain of *Ftm*^*-/-*^ embryos at E8.5. At E12.5, cilia were severely reduced in number, and their shape and content were highly abnormal. This, combined with our previous studies (Besse et al., 2011), strongly suggests that the forebrain defects observed in *Ftm* mutants are due to the ciliary defects in neural progenitors.

Our data point to region-specific defects in Hh/Gli signaling in the forebrain of *Ftm* mutants. The reduction in ventral forebrain areas and the loss of the TH-R in *Ftm* mutants suggest an impaired response to Shh signals, similar to what has been previously observed in the ventral spinal cord of these mutants (Vierkotten et al., 2007). Indeed, tegmental areas of the diencephalon and hypothalamus depend on Hh signaling from the notochord and prechordal plate, which induces *Shh* expression in the forebrain midline and neural Shh is in turn required for correct formation of the basal diencephalon and hypothalamus (Dale et al., 1997; Szabo et al., 2009a; 2009b; Shimogori et al., 2010; Zhao et al., 2012). High Hh activity (from the ZLI and the ventral forebrain) is also required for the formation of the TH-R, while the TH-C requires lower Hh activity (Hashimoto-Torii et al., 2003; Jeong et al., 2011). Thus, our observation of the loss of the TH-R and the expansion of the TH-C in *Ftm*^*-/-*^ embryos is totally consistent with the strong reduction of GliA activity as assayed by Tg[GBS::GFP] and the near-total absence of *Shh* expression in the ventral diencephalon.

However, the phenotype of the *Ftm* mutant in the forebrain differs from that of a *Shh* mutant. In *Shh*^-/-^ embryos, unlike in *Ftm*^*-/-*^ embryos, the whole diencephalon is extremely reduced in size due to reduced proliferation and survival as soon as the 15s stage (Ishibashi and McMahon 2002). Moreover, contrary to *Shh* mutants (Chiang et al., 1996), *Ftm* mutants never show cyclopia, even when two copies of Gli3R are reintroduced. This suggests that a low level of Hh pathway activity (undetected by the GBS::GFP transgene) sufficient to separate the eye fields and to promote forebrain morphogenesis is produced from the underlying prechordal mesendoderm of *Ftm*^*-/-*^ embryos.

In that respect, examination of Hh/Gli activity at the ZLI is very informative. Indeed, the ZLI forms in *Ftm*^-/-^ embryos and it is even wider than in controls. This widening is accounted for by the reduction in Gli3R levels, since it is rescued in compound [*Ftm*^*-/-*^*, Gli3*^*Δ/+*^] embryos. Consistent with this data, Gli3 repression by Wnt signals is required for controlling the width of the ZLI in chicken embryos (Martinez-Ferre et al., 2013). Moreover, *Shh* from the ZLI appears to be able to signal, although with lower efficiency than in controls. Thus, the Hh/Gli signaling pathway is still active in *Ftm*^-/-^ embryos. Moreover, a basal, low level of Gli activity appears to be present throughout the diencephalon, likely caused by the reduction in Gli3R levels (Besse et al., 2011).

The ZLI has been proposed, initially in chick, to form through an inductive process requiring Hh signaling from the diencephalic basal plate (Kiecker and Lumsden, 2004; Zeltser, 2005, Epstein, 2012). In mouse mutants in which expression of a functional Shh is absent from the ventral diencephalon, the ZLI does not form (Szabo et al., 2009b). If ZLI formation requires Hh signals from the basal plate, how can it occur in *Ftm* mutants, which display no *Shh* expression in the basal diencephalon? In E9.5 *Ftm*^-/-^ embryos, a discrete patch of *Shh* expression remained in the basal plate at the level of the future ZLI. We propose that this patch of *Shh* expression is sufficient for the initiation of ZLI formation in *Ftm* mutants. Because this patch does not give rise to detectable Gli activity, we speculate that here Shh might signal through Gli-independent, non-canonical pathway (Carballo et al., 2018).

Examination of the eye in *Ftm* mutants provides another example of the region-specific functions of cilia. In *Ftm*^*-/-*^ embryos, the optic cup and lens are totally absent. Gli3 is known to be involved in optic cup formation (Furimsky and Wallace, 2006), but it was not known so far whether it acted as a repressor or as an activator. We found optic cups with correct DV patterning in compound [*Ftm*^*-/-*^*, Gli3Δ/+*] *and* [*Ftm*^*-/-*^*, Gli3Δ/Δ*] embryos, showing that Gli3R, and not Gli3A, is crucial for optic cup formation, and that the function of cilia in this process is mediated by Gli3R. This was confirmed by the analysis of [*Ftm*^*+/+*^*, Gli3Δ/Δ*] siblings, which displayed normal retina. The retinal phenotype of *Ftm* mutants is reminiscent to that of the telencephalon, where dorsal structures are reduced due to the reduction in Gli3R levels (Besse et al. 2011; Laclef et al., 2015). However, in compound [*Ftm*^*-/-*^*, Gli3Δ/+*] and [*Ftm*^*-/-*^*, Gli3Δ/Δ*] embryos, the optic cups were closer to each other under the ventral forebrain and even partially fused in some cases, and the optic stalk was almost totally absent. Thus, cilia are required both for GliR-dependent optic cup formation and for GliA-dependent optic stalk morphogenesis.

In conclusion, our data show that, in *Ftm* mutants, forebrain structures requiring high GliA activity, such as the rostral thalamus and ventral forebrain, and structures that require high GliR activity, such as the optic cup, are lost. In contrast, structures that require low or intermediate Hh activity, such as TH-C or the optic stalk, are still present. Thus, different regions of the forebrain are differently affected by the loss of cilia depending on their specific requirement for GliA or GliR activity (Figure 10B-D).

Are our data relevant for human disease? There are few reports of hypothalamic or diencephalic malformations in ciliopathies. However, a precise analysis of the forebrain is rarely possible in fetuses with severe ciliopathies such as Meckel syndrome. Microphthalmia and benign tumors called diencephalic or hypothalamic hamartomas have been observed in Meckel syndrome and other ciliopathies (Ahdab-Barmada and Claassen, 1990; Roume et al.,1998; Paetau et al., 2008; Poretti et al. 2011; Del Giudice et al., 2014; Poretti et al., 2017). Interestingly, diencephalic hamartomas have been linked to mutations in *GLI3* and other SHH pathway genes (Shin et al., 1999; Hildebrand et al., 2016), suggesting that those observed in ciliopathies could also be caused by defects in SHH signaling. Holoprosencephaly is rarely described in ciliopathies, and only in the most severe form, Meckel syndrome (Paetau et al., 1985, Ahdab-Barmada and Claassen, 1990). This may be surprising, given the essential role of cilia in vertebrate Hh signaling. Our study of the forebrain of *Ftm* mutants provides a potential explanation, as we find clear phenotypic differences between the *Ftm* mutants and Hh pathway mutants. Nevertheless, ciliopathy genes could act as modifier genes for HPE. HPE shows high phenotypic variability in single families, which has led to the proposal that a combination of mutations in HPE genes could account for the variable severity of the phenotype (the multi-hit hypothesis). In favor of this hypothesis, digenic inheritance has been identified in several HPE families (Mouden et al., 2016 and ref therein). Interestingly, homozygous mutations in the *STIL* gene encoding a pericentriolar and centrosomal protein have been found in patients with HPE and microcephaly (Mouden et al., 2015, Kakar et al., 2015). Mouse *Stil*^-/-^ embryos display severe forebrain midline defects (Izraeli et al., 1999) and ciliogenesis, centriole duplication and Hh signaling are defective in the absence of STIL (David et al., 2014; Mouden et al., 2015). Whole genome sequencing in heterogenous HPE families will allow testing the involvement of ciliopathy gene variants in this disease.

More generally, our study of the ciliopathy gene mutant *Rpgrip1l/Ftm* calls for further examination of ciliary and ciliopathy genes in human neurodevelopmental diseases associated with SHH pathway defects.

## Acknowledgements

We are grateful to the animal and imaging facilities of the IBPS (Institut de Biologie Paris-Seine FR3631, Sorbonne Université, CNRS, Paris, France) for their technical assistance. We thank Michaël Trichet (electron microscopy facility, IBPS) for electron microscopy analysis. We are grateful to Christine Vesque and Marie Breau (IBPS-Developmental Biology laboratory, Paris, France) and Christoph Gerhardt (Institute for Animal Developmental and Molecular Biology, Heinrich Heine University Düsseldorf, Germany) for critical reading of the manuscript. We thank James Briscoe (Crick Institute, London, UK) for the kind gift of the Tg[GBS::GFP] transgenic line. This work was supported by funding from the Agence Nationale pour la Recherche (ANR, project CILIAINTHEBRAIN to SSM), the Fondation pour la Recherche Médicale (Equipe FRM DEQ20140329544 funding to SSM) and the Fondation ARC pour la Recherche sur le Cancer (Project ARC PJA 20171206591 to SSM).

